# Spinal and cortical premotor control of primate dexterous hand movements

**DOI:** 10.1101/2024.09.14.613086

**Authors:** Tomohiko Takei, Tomomichi Oya, Kazuhiko Seki

## Abstract

Skilled hand movements are an evolutionary advance in primates. Phylogenetically distinct corticospinal pathways are involved in hand control: “newer” direct corticomotoneuronal (CM) pathways and “older” indirect corticospinal pathways mediated by spinal premotor interneurons (PreM-INs). However, the functional differences of these pathways remain unclear. Here we show that CM cells and PreM-INs have distinct physiological properties for the activation of hand muscles and provide different types of movement control signals while monkeys perform a precision grip task. Spike-triggered averaging of electromyographic activity indicated that PreM-INs coactivate a larger number of muscles, whereas CM cells more selectively control fewer muscles. The firing activity of PreM-INs was tightly correlated with their target muscle activity and had a greater contribution to generating hand muscle activity. In contrast, CM cell activity diverged temporally from the target muscle activity and had a smaller contribution to its generation. On the basis of these results, we hypothesize that PreM-INs produce gross muscle activity by activating synergistic muscles, whereas CM cells fine-tune target muscle activity. This idea was supported by dimensional reduction analyses of hand muscle activity, as PreM-IN activity was specifically correlated with lower dimensional control of muscle activity, and CM cell activity was correlated with higher dimensional control. These results indicate that the two pathways have distinct functions, synergistic control and fine tuning of hand muscle activity, both of which are essential for the development of dexterous hand movement in primates.

## Introduction

Physiological and anatomical studies have suggested that phylogenetically distinct pathways are involved in the control of dexterous hand movements in primates. A direct corticomotoneuronal (CM) pathway is a phylogenetically newer pathway that has evolved specifically in primates^1–4^. The development of CM cells correlates with the development of dexterous hand function in primates, thus it is believed to play a major role in the control of dexterous hand movements^1,2^.

In contrast, indirect corticospinal pathways, mediated by spinal premotor interneurons (PreM-INs), represent phylogenetically older pathways commonly observed in a diverse range of mammals^1,2^. As these pathways are present in species with less hand dexterity^1^, their contribution to the control of skilled hand movements has been less appreciated. However, our recent studies demonstrated that spinal PreM-INs are involved in controlling skilled hand movements in primates. These studies showed that spinal PreM-INs provided post-spike facilitation to the hand muscle during a precision grip^5^. In addition, their activity was correlated with the target muscle activity and grip force^6,7^. These findings suggest that the older PreM-IN pathways also contribute to the control of skilled hand movements in primates, alongside the CM cells.

These findings raise the question of how the PreM-IN and CM systems concurrently contribute to dexterous hand movements in primates, and whether they provide structural redundancy in the nervous system or fulfil distinct functions. Here, we hypothesized that the two systems have different roles and that their cooperative actions are essential for the evolution of dexterous hand movements in primates.

To test this hypothesis, we compared how PreM-INs and CM cells contribute to the generation of hand muscle activity by analysing spike-triggered averages of muscle activity during a precision grip task. Our findings showed that a PreM-IN exhibited post-spike facilitation on a larger number of hand muscles, and its firing activity had a higher correlation with the target muscle activity than was measured with a CM cell. Through the reconstruction of electromyographic (EMG) activity generated by a single PreM-IN or CM cell, computed by convolving post-spike effects with firing activity, we demonstrated that PreM-INs have a greater contribution than CM cells to the generation of hand muscle activity during the precision grip task. Furthermore, dimensional reduction analyses (muscle synergy model) demonstrated that PreM-IN activity was correlated with lower dimensional control of hand muscles, whereas CM cell activity was correlated with the higher dimensional components of target muscle activity.

These results suggest that the two pathways have distinct roles and that their cooperation is essential for skilled hand movement in primates. Specifically, PreM-INs produce a gross pattern of muscle activity through the integration of synergistic muscle activity. In contrast, CM cells finely tune individual muscle activity. The coexistence of synergistic and flexible control systems may underlie the evolution of dexterous hand function in primates.

## Results

### PreM-INs coactivate a larger group of muscles whereas CM cells facilitate fewer muscles

To test the difference in the output effects of PreM-INs and CM cells on hand muscle activity, we examined the post-spike facilitation of PreM-INs and CM cells. We recorded single-unit activity from the cervical spinal cord or hand region of the primary motor cortex in two monkeys performing a precision grip task (Extended Data Fig. 1). Using a spike-triggered average of EMGs, we identified 14 PreM-INs and 19 CM cells that showed post-spike facilitation in hand and arm muscles. Because of the small sample size, we excluded neurons with post-spike suppression from our analysis (five PreM-INs and two CM cells). Figure 1a shows examples of post-spike facilitation produced by PreM-INs and CM cells. A substantial population of neurons in both groups exhibited post-spike facilitation on more than one hand muscle, commonly referred to as the muscle field^8,9^ (Extended Data Fig. 2 for all neuron data). We compared the size of the muscle field, defined as the number of muscles facilitated by each neuron, and found that the muscle field was larger for PreM-INs than for CM cells (3.1 ± 2.3 vs 1.8 ± 1.1, *P* < 0.05, *t*-test, *t*_[31]_ = 2.16, Fig. 1b, Extended Data Fig. 4a for each animal), indicating that PreM-INs coactivated a larger number of hand muscles. In contrast, CM cells facilitated a smaller number of muscles, and more than half of them (12/19 cells) had a post-spike facilitation on a single muscle.

**Figure 1.**
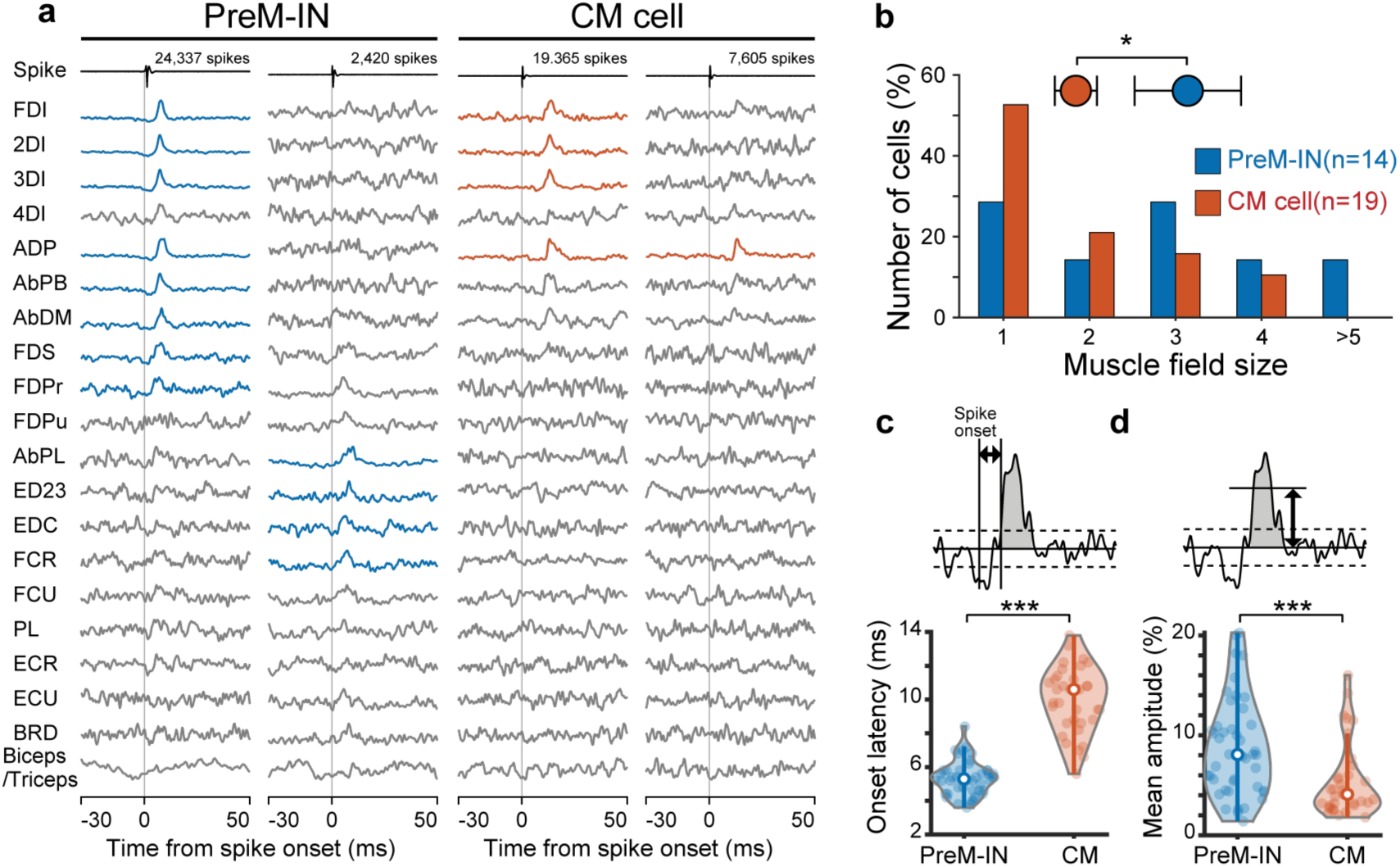
Comparison of post-spike facilitations of PreM-INs and CM cells. **a**, Spike-triggered averages of rectified electromyography (EMG) signals by example premotor interneurons (PreM-INs, left) and corticomotoneuronal cells (CM cells, right). Spike-triggered averages with significant post-spike facilitations are indicated in blue (PreM-IN) or red (CM cell). **b,** Muscle field size. Circles, means. Error bars, standard error of means. \**t*-test, *P* < 0.05. **c–d**, Comparison of onset latency (**c**) and mean amplitude increase (**d**). A filled dot indicates a post-spike effect. Horizontal bars, means. *** *t*-test, *P <* 0.001.

Next, we compared the onset latency and size of the post-spike facilitation between the groups (Fig. 1c–d, Extended Data Fig. 4b–c). The onset latency was shorter for PreM-INs than for CM cells (5.3 ± 1.0 vs 10.0 ± 1.9 ms, *P* < 0.001, *t*-test, *t*_[48.85]_ = −12.85), which was expected due to the relative proximity of muscles to the spinal cord compared with the cerebral cortex. In addition, the post-spike facilitation size was larger in PreM-INs than in CM cells (8.8% ± 4.9% vs 5.4% ± 3.7%, *P* < 0.001, *t*-test, *t*_[76.79]_ = 3.49), particularly in one monkey (monkey S, Extended Data Fig. 4c). These results indicate that PreM-INs coactivate a larger number of muscles with a shorter latency and greater amplitude, whereas CM cells have post-spike effects on a limited set of muscles with a longer latency and smaller amplitude.

### PreM-IN has target muscle-like activity, whereas a CM cell diverges temporally from target muscles

Given the faster and stronger facilitation observed in PreM-INs to the target muscles, we anticipated that PreM-IN activity would exhibit a closer temporal relationship with the target muscle activity than would be observed with CM cells. To test this, we computed the cross-correlation between the neural firing activity and the target muscle activity. Figure 2a–d shows examples of the cross-correlations between the activity of a PreM-IN and CM cell with their target muscle activity. The cross-correlogram demonstrated that the PreM-IN activity had a high correlation value (*Rmax* = 0.84) with a peak lag of −4.8 ms, indicating that the PreM-IN activity preceded the target muscle activity with a short latency (Fig. 2b). The cross-correlation of a CM cell showed a lower correlation value (*Rmax* = 0.22), and it did not exhibit a clear peak preceding the target muscle activity (Fig. 2d). Figure 2e–f shows the distribution of *Rmax* and peak lag for neuron–muscle pairs of PreM-INs and CM cells. On average, the PreM-INs had a higher correlation value (0.46 ± 0.19), with the peak lag clustered at around zero (4.6 ± 0.14 ms), whereas CM cells had a lower correlation value (0.26 ± 0.25, *P* < 0.001, *t*-test, *t*_[62.11]_ = 3.86, Fig. 2g, Extended Data Fig. 4d), and the lag was broadly distributed (−28.5 ± 274.6 ms, *F*-test for equal variances, *P* < 0.001, *F*_[43, 34]_ = 0.26, Fig. 2h, Extended Data Fig. 4e). These results suggest that PreM-INs have more target muscle-like activity, whereas CM cells diverge temporally from the target muscle activity.

**Figure 2.**
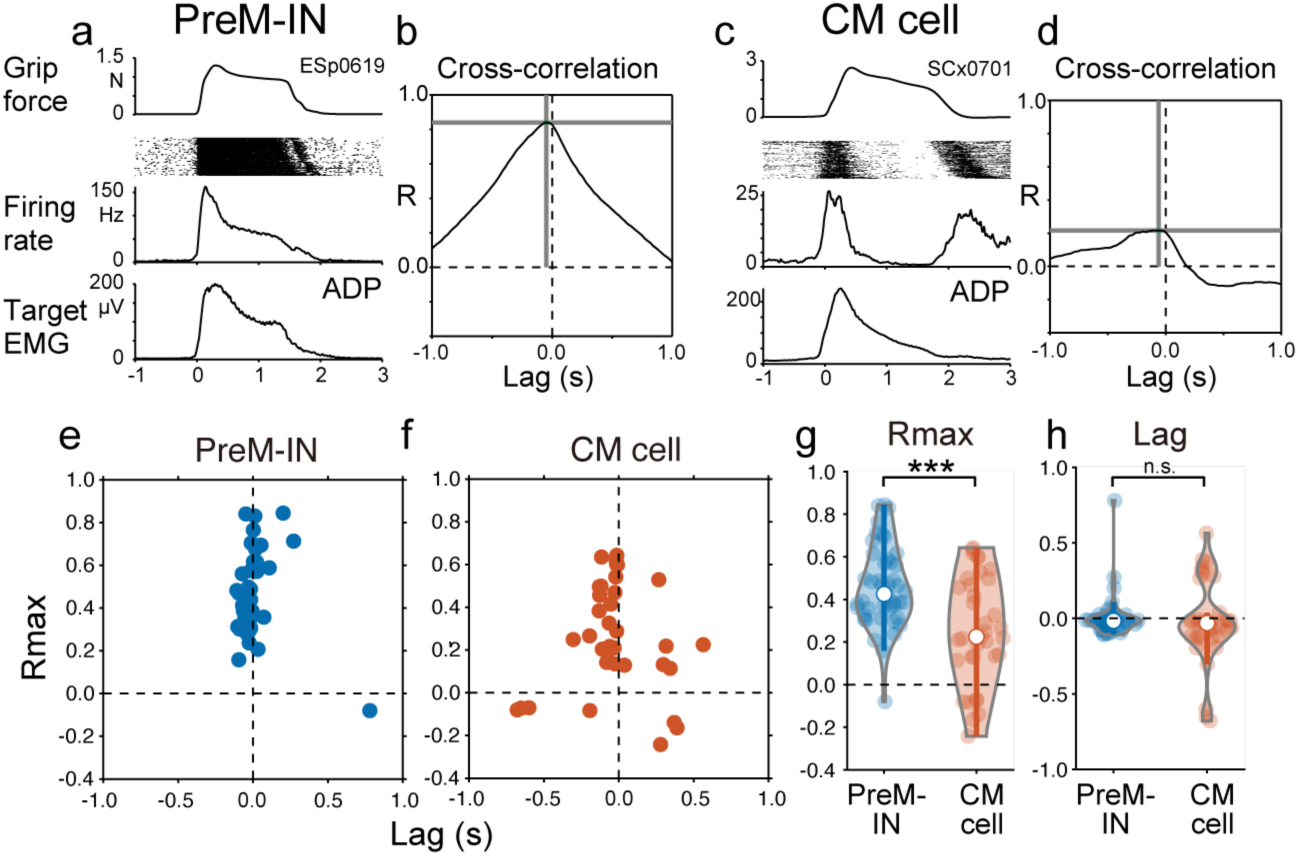
Temporal correlation with target muscle activity. **a–b**, Activity profile of an example PreM-IN and one of its target muscles (adductor pollicis, ADP). **b**, cross-correlogram between them. The peak correlation value (*Rmax*) and its lag is indicated by horizontal and vertical grey lines. c–d, Same format but for an example CM cell and target muscle pair. e–f, Distribution of *Rmax* and lag for PreM-INs (**e**) and CM cells (**f**). g–h, Comparison of *Rmax* (**g**) and lag (**h**). A filled dot indicates a neuron–muscle pair. Horizontal bars, means. *** *t*-test, *P <* 0.001. n.s. non-significant.

### PreM-INs have a greater contribution to generation of hand muscle activity

To assess the contribution of PreM-INs and CM cells in the generation of hand muscle activity, we estimated the EMG activity generated by individual neurons and compared their relative amplitude to the original EMG activity. We reconstructed EMG activity produced by a single neuron by convolving a waveform of post-spike facilitation with the firing activity of the triggering neuron (Fig. 3a). Examples of reconstructed EMG of a single PreM-IN and CM cell are shown in Figure 3b and c. The reconstructed EMG of a PreM-IN had a temporal profile similar to that of the original EMG, reflecting the high temporal correlation between the neurons and target muscles (Fig. 3b). To quantify the relative contribution of individual neurons, we calculated a ratio of the area under the EMG profiles during a grip period (reconstructed/original, %). The example PreM-IN contributed 1.9% (5.7/298.9 µV*s) to the original EMG activity, whereas the reconstructed EMG by a CM cell had a smaller contribution of 0.6% (1.3/240.6 µV*s) (Fig. 3c). On average, PreM-INs contributed approximately four times more to target muscle activity generation than CM cells (1.01% ± 0.73% vs 0.25% ± 0.68%, *P* < 0.001, *t*-test, *t*_[74.81]_ = 4.75, Fig. 3d, Extended Data Fig. 4f).

**Figure 3.**
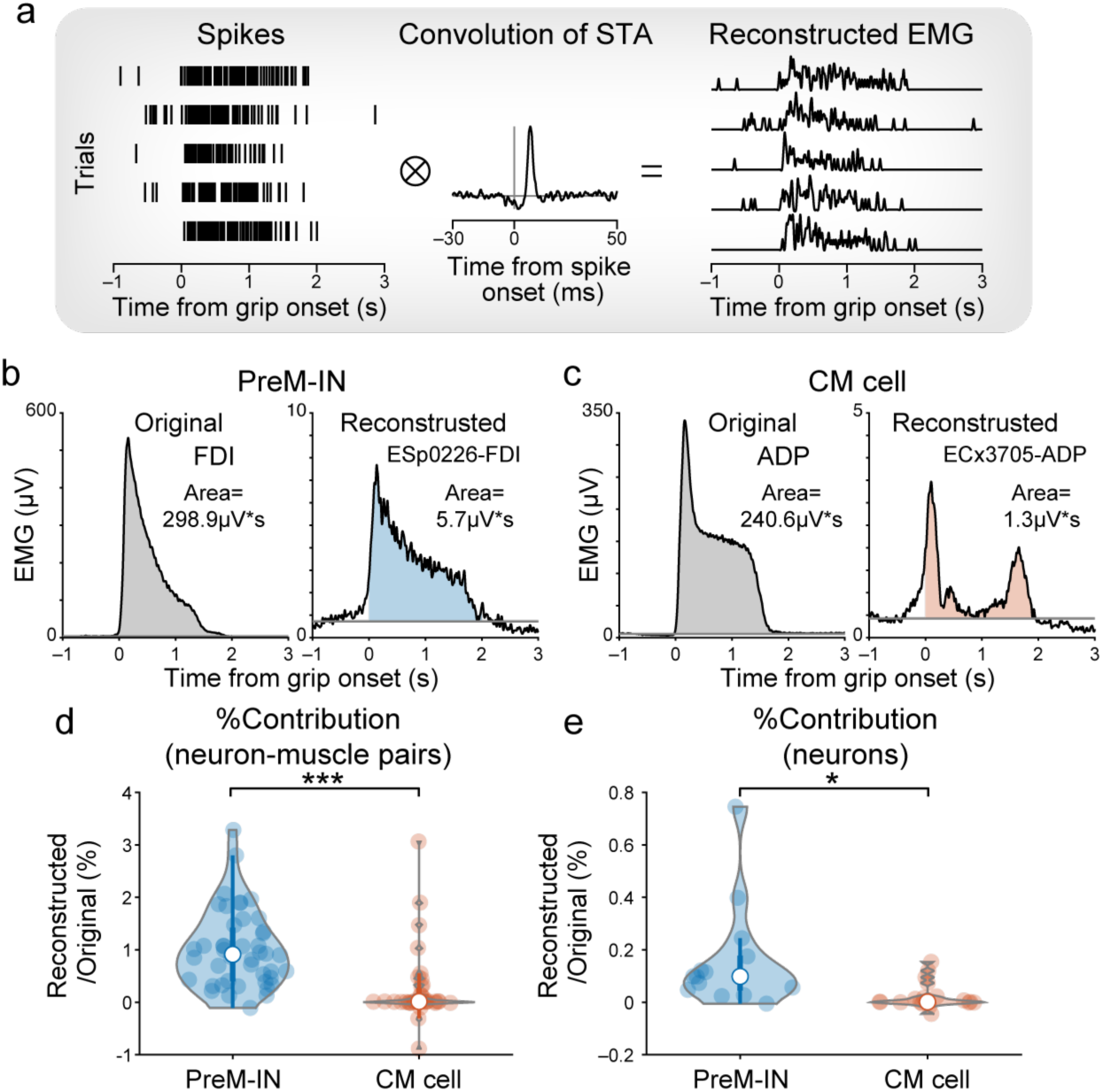
Reconstruction of EMG from single neuron activity. **a**, Muscle activity generated by a single neuron was estimated by convolving spike firing and a waveform of a post-spike effect. **b**–**c**, Example of muscle output by a single PreM-IN (**b**) and CM cell (**c**). **d–e**, Relative contribution of single neuron activity to generation of a single target muscle activity (**d**) and entire recorded muscles (**e**). A filled dot indicates a neuron–muscle pair (**d**) or a neuron (**e**). Horizontal bars, means. * *t*-test, *P <* 0.05. *** *P* < 0.001

Given that PreM-INs had a larger muscle field (Fig. 1), their contribution to generating the activity of individual muscles could be magnified when considering the activity of all muscles required for performing the task. To investigate this, we calculated the relative contribution to all recorded muscles for each individual neuron (Fig. 3e). We found that the relative contribution of PreM-INs (0.16% ± 0.20%) was eight times greater than that of CM cells (0.02% ± 0.05%, *P* < 0.05, *t*-test, *t*_[14.24]_ = 2.49, Fig. 3e, Extended Data Fig. 4g). These results suggest that a PreM-IN contributes to the formation of gross activity over multiple muscles during precision grip, whereas a CM cell provides fine-tuning of the muscle activity in a relatively individual manner.

### PreM-INs and CM cells involved in lower and higher dimensional control, respectively, of hand muscle activity

Our results demonstrated distinct properties of PreM-INs and CM cells for controlling hand muscle activity during a precision grip task. Considering these observations, we formulated a hypothesis that the two pathways serve different functions in the control of hand muscles: PreM-INs are responsible for generating gross muscle activity by synergistically activating multiple muscles, whereas CM cells fine tune the activity of individual target muscles. To directly test this hypothesis, we compared the PreM-IN and CM cell activity with the muscle synergy model^10^. The muscle synergy model assumes that a hand muscle’s activity is generated through a linear combination of a small number of low dimensional modules referred to as muscle synergies. Conventionally, muscle synergies explain a majority (approximately 90%) of the variance of the overall hand muscle activity^10^. However, it is important to note that a considerable portion of variance (approximately 10%), referred to as the “residuals”, remains unexplained and is often ignored as noise. Recent studies, however, highlighted the unexpected complexity of primate hand movements and suggested the higher dimensional variance, not captured by low dimensional components, may incorporate significant volitional control^11–13^. Therefore, we modelled hand muscle activity during a precision grip as a combination of lower-dimensional control by muscle synergies and higher-dimensional control represented in the residuals (Fig. 4a). Our hypothesis predicted that the PreM-IN activity would correlate with the muscle synergy, whereas the CM cell activity would correlate with the residual of the target muscle activity.

**Figure 4.**
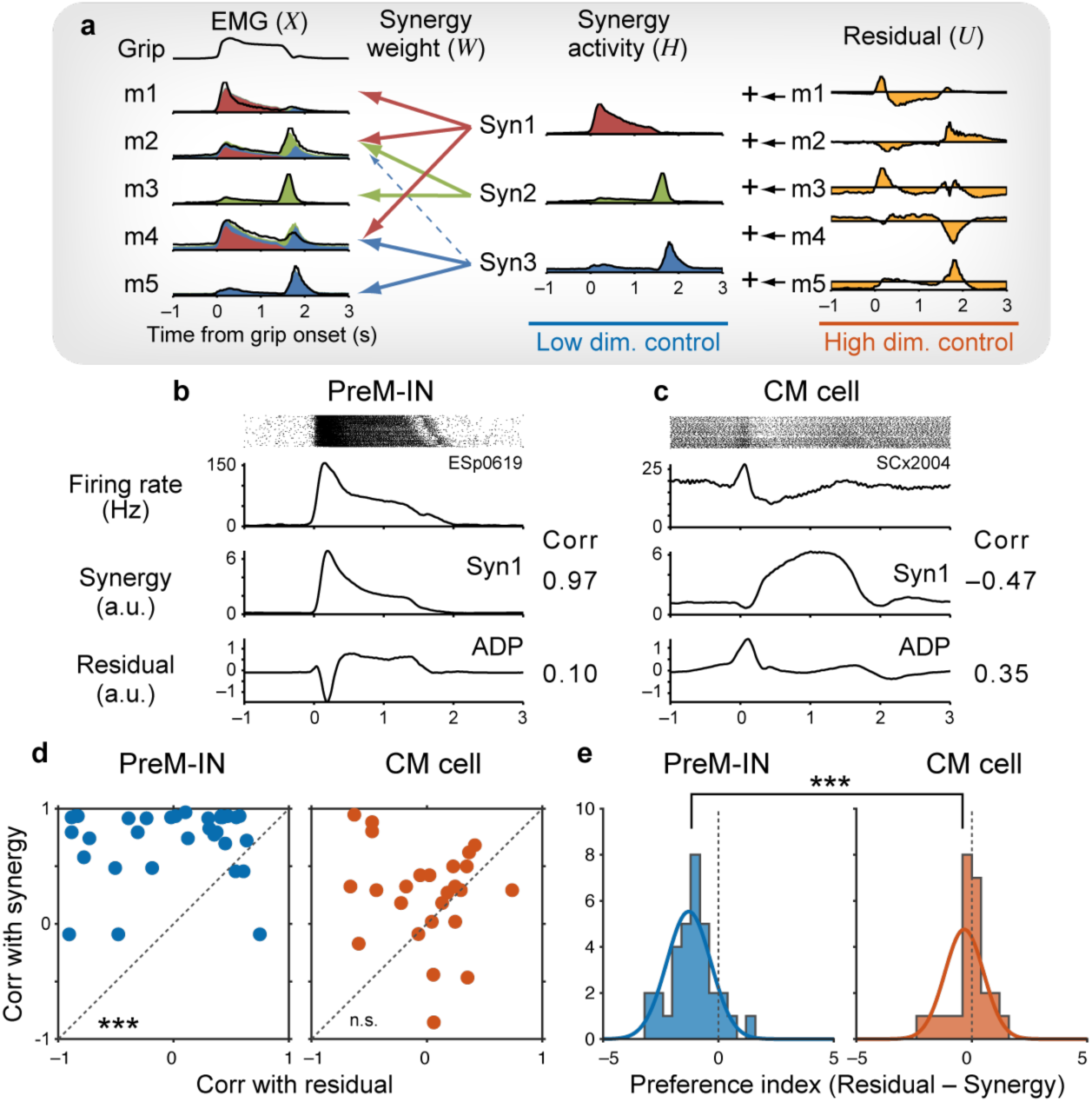
Comparison with a muscle synergy model. **a**, Muscle synergy model. A linear combination of a small number of muscle synergies explain majority (approximately 90%) of variance of original EMG, and residuals (approximately 10%) is added to the individual muscles. **b–c**, Example firing activity of PreM-IN and CM cells, and a preferred synergy and a target muscle activity. **d**, Correlation of PreM-INs (left) or CM cell activity (right) with the preferred synergy and residual of the target muscle activity. **e**, Preference index of PreM-INs (left) or CM cells (right). *** *t*-test, *P <* 0.001. n.s. non-significant

Dimensional reduction analysis showed that a small number of muscle synergies (three and four in monkeys E and S, respectively) explained approximately 90% of the variance in muscle activity (Extended Data Fig. 3). Figure 4b shows an example of the correlation between PreM-IN activity and the preferred muscle synergy, whose synergy weights showed the highest similarity with the neuron’s muscle field^10^, and with residual of one of the target muscles. This PreM-IN showed a high correlation with the preferred synergy and low correlation with the target muscle residual (Fig. 4b). In contrast, an example CM cell showed a higher correlation with the residual and lower correlation with the preferred synergy (Fig. 4c). Figure 4d provides a comparison of the correlations between each neuron’s activity with the preferred synergy and the residual of the target muscle activity. On average, a PreM-IN had a higher correlation with synergy than with residual (0.73 ± 0.31 vs −0.04 ± 0.56, *P* < 0.001, paired *t*-test with *z* transformation, *t*_[32]_ = 7.76, Fig. 4d, Extended Data Fig. 4h) whereas a CM cell was more evenly correlated with both the synergy and residual (0.25 ± 0.43 vs −0.00 ± 0.37, *P* = 0.063, paired *t*-test with *z* transformation, *t*_[24]_ = 1.95, Fig. 3d, Extended Data Fig. 4i–j). To assess the preference of each group of neurons, we computed a preference index (Fig. 4e) and found that PreM-IN showed a significantly higher preference for synergies compared with residual, whereas the CM cell had a closer preference for the synergy and residual (−1.28 ± 0.95 vs −0.33 ± 0.84, *P* < 0.001, *t*-test, *t*_[54.67]_ = −4.07, Fig. 4e, Extended Data Fig. 4k–m). These results supported our hypothesis that PreM-INs play a role in the lower-dimensional control necessary for generating gross hand muscle activity, whereas the CM cells add higher-dimensional control essential for providing flexibility of the target muscles’ activity.

## Discussion

In this study, we demonstrated distinct properties of PreM-IN and CM cells in the control of skilled hand movements in primates. PreM-INs exhibited a synergistic coactivation of a larger number of muscles, suggesting the generation of gross hand muscle activity. In contrast, CM cells showed more selective control of individual muscles, suggesting fine-tuning of target muscles. The low dimensional control of muscles via muscle synergies, as observed in PreM-INs, has been previously associated with enhancing the efficiency and robustness of motor control^14–19^. In contrast, fine-tuning of individual muscle activity is essential for achieving fractionated control of hand muscles^9,20^ and relatively independent finger movements^21,22^. Our results provide direct physiological evidence that these two aspects of hand motor control are mediated by distinct premotoneuronal systems with different properties. Given that both systems come into play during skilled hand movement, such as precision grip, it is conceivable that the coordination of the efficient (synergistic) and flexible (fractionated) control systems, rather than one or the other in isolation, is essential for achieving the remarkable hand dexterity in primates.

Over the past 40 years, studies have consistently demonstrated the significant role of CM cells in voluntary control of skilled hand movements, including precision grip^9,20,23–25^, relatively independent finger movements^26–28^, and wrist movements^8,29,30^. Notably, these studies have demonstrated that CM activity does not simply reflect target muscle activity. For example, Muir and Lemon showed that a CM cell was selectively activated during a precision grip but was deactivated during a power grip, even though the target muscle remained active^23^. Similarly, Griffin et al. showed that the relationship between CM cells and target muscles was broadly distributed^29^. Our findings suggest that this unexplained nature of CM cells reflects their contribution to high-dimensional control of hand muscles, which was not captured by the low-dimensional control responsible for a majority of variance (Fig. 4a). Given that CM cells have a smaller muscle field (Fig. 1b) and make a smaller contribution to the generation of muscle activity (Fig. 3d–e), it is plausible that their role is to provide fine-tuning of target muscles on top of the lower-dimensional synergistic control. In contrast, the phylogenetically older PreM-INs play a more substantial role in generating gross hand muscle activity (Fig. 3d–e) and are specifically correlated with the lower-dimensional control (Fig. 4a). These results suggest that over the course of primate evolution, the CM system did not replace or override the function of the older systems^2^, but rather added a newer layer of control that enhances the flexibility of primate hand motor control.

It is important to note that although there is a relative difference between the CM cells and PreM-INs, there is also a large overlap in their functional properties. This overlap is evident in various aspects, including the distribution of muscle field size (Fig. 1b), temporal correlation with target muscle activity (Fig 2g), and the preference index of muscle synergy and residuals (Fig. 4e) between the groups. These observations suggest that the functional differentiation of these two systems is not mutually exclusive but rather is gradual and characterized by significant overlap. During neonatal development, it has been demonstrated that corticospinal terminals gradually expand from the intermediate zone to the ventral horn over the first 5 months in the first thoracic segment of monkey spinal cord^31^. This anatomical shift suggests that the corticospinal tract progressively incorporates the indirect connections and direct CM projections, leading to the gradual overlap of functional properties between the two systems.

A previous comparison of CM cells and PreM-INs in the context of voluntary wrist movements demonstrated that spinal PreM-INs have a smaller muscle field than the CM cell^32^, which conflicts with our present findings. Several possible reasons might account for this discrepancy. First, wrist movements typically involve the reciprocal activation of antagonist muscles, whereas hand grasping often requires the coactivation of antagonistic muscles^33–35^. Notably, during wrist movements, a significant number of CM cells showed a reciprocal muscle field, characterized by both excitatory and inhibitory post-spike effects on antagonistic muscles at the same time^36^, a phenomenon not observed in our present study. This suggests that different control strategies are used during wrist movements (reciprocal) and grasping (coactivating). Second, the previous study compared CM cells and PreM-INs based on the data compiled from different experiments, which used different criteria to identify the post-spike effects^8,32,37^. In the present study, we used consistent analytical procedures for both cortical and spinal recordings in the same animals, enabling us to directly compare the detailed physiological properties.

Spinal PreM-INs receive convergent inputs from various descending tracts and peripheral afferents^38^. This indicates that the lower-dimensional control mediated by the spinal PreM-INs may serve as a fundamental platform for both voluntary and reflexive hand functions in primates. Interestingly, these pathways also have unique connections to directly control motoneurons^37^. Further research aimed at identifying the descending and sensory inputs to PreM-INs, and comparing them with the direct control of hand motoneurons, would provide new insights into the functional divergence in the descending and spinal control of skilled hand movements.

## Methods

### Dataset

The datasets used in the present study were obtained from two male macaque monkeys (monkey E: *Macaca mulatta*, 5.6 kg at the age of 5 and monkey S: *Macaca fuscata*, 9.0 kg at the age of 8). The spinal recordings dataset from monkey E was the subject of previous reports^1–4^. The rest of the datasets (cortical recordings from monkeys E and S and spinal recordings from monkey S) were newly collected for this study. Only a post-spike suppression was identified in the spinal recording of monkey S, which was excluded from the present report. Therefore, all spinal data presented in this paper were obtained from monkey E. All procedures were approved by the Animal Research Committee at the National Institute for Physiological Sciences, Aichi, Japan and the experimental animal committee of the National Institute of Neuroscience, Tokyo, Japan.

### Behavioural task

The monkeys were trained to grip spring-loaded levers with their left index finger and thumb (precision grip task, Extended Data Fig. 1a)^1,5^. Lever positions were displayed on a computer screen as cursors and monkeys were required to track targets in a step-tracking manner. Each trial comprised a rest period (1.0–2.0 s), lever grip, lever hold (1.0–2.0 s), and lever release. Successful completion of a trial was rewarded with a drop of apple sauce (Extended Data Fig. 1d). The force required to reach the target positions was adjusted independently for the index finger and thumb of each monkey.

### Surgical procedures

After the task training was completed, we performed surgeries to implant head restraints, EMG wires, and recording chambers under isoflurane or sevoflurane anaesthesia and aseptic conditions. For EMG recordings, we subcutaneously implanted pairs of stainless steel wires (AS 631, Cooner Wire, Chatsworth CA, USA) in the forelimb muscles, including intrinsic hand muscles (first, second, third and fourth dorsal interosseous, FDI, 2DI, 3DI and 4DI; adductor pollicis, ADP; abductor pollicis brevis, AbPB; abductor digiti minimi, AbDM), extrinsic hand flexors (flexor digitorum superficialis, FDS; radial and ulnar parts of the flexor digitorum profundus, FDPr and FDPu), wrist flexors (flexor carpi radialis, FCR; flexor carpi ulnaris, FCU), extrinsic hand extensors (extensor digitorum-2,3, ED23; extensor digitorum communis, EDC), a wrist extensor (extensor carpi ulnaris, ECU), elbow muscles (brachioradialis, BRD; biceps brachii, Biceps).

For spinal recordings, we implanted a recording chamber on the cervical vertebrae (C4– C7) of monkeys, where a unilateral laminectomy was made on the ipsilateral side of the employed hand (Extended Data Fig. 1b).

For cortical recordings, we implanted a recording chamber (a circular cylinder with a 50-mm diameter) over a craniotomy covering a cortical area, including the hand representation of pre– and post-central gyri on the contralateral side of the employed hand (Extended Data Fig. 1c).

### Data recordings

While the monkey was performing the precision grip task, we recorded single-unit activity from cervical 6 (C6)–Thoracic 1 (T1) segments or from the hand area of the primary motor cortex with a tungsten or Elgiloy microelectrode (impedance 1–2 MΩ at 1 kHz) with a hydraulic Microdrive (MO-951, Narishige Scientific Instrument, Tokyo, Japan). We determined the recording sites with the aid of positions of vertebral segments for the spinal recordings, and the sulcal structures and intercortical electrical microstimulation (ICMS, 300 Hz, 10 pulses, <20 µA) for the cortical recordings. Action potential timing was detected online using a spike-sorting device (MSD; Alpha Omega Engineering, Nof HaGalil, Israel), and spike-triggered averaging of rectified EMG signals was monitored during recording. Neural signals and spike times were digitized at 25 kHz. EMGs were bandpass-filtered (5 Hz to 3 kHz) and sampled at 5 kHz. Simultaneously with the neural and EMG signals, we recorded grip force and behavioural event timing at 1 kHz (Extended Data Fig. 1e).

### Identification of post-spike effects of spinal PreM-INs and CM cells

Detailed analyses of the post-spike effects of spinal neurons^1^ and their activity^2,3^ were reported previously. All analyses were performed off-line using MATLAB (MathWorks). The spike-triggered average of the rectified EMG was computed to identify post-spike effects of the spinal or cortical neurons on the recorded EMGs^1–4^. We analysed only neurons with ≥2000 recorded spikes. Spike-triggered averages were compiled by averaging segments of rectified EMG activity from 30 ms before to 50 ms after each trigger. Spikes were accepted as triggers only if the root mean square (RMS) value of the EMG from 30 ms before to 50 ms after the spike was greater than 1.25 times the RMS noise level in that EMG channel. The spike-triggered average was smoothed with a flat five-point finite impulse response filter. The baseline trend was subtracted using the incremented-shifted averages method^6^, and significant spike-triggered average effects were identified with multiple-fragment statistical analysis^7^. The test window was set to 3–15 ms after the spinal neuron spike and 6–16 ms after the cortical neuron spike^8^.

Potential cross-talk between recorded EMGs was evaluated by combining a cross-correlation method^9^ and the third-order differentiation^10^, and spike-triggered average effects potentially resulting from cross-talk between EMG recordings were eliminated from the present data set. To distinguish the post-spike effects from the synchrony effects^8^, we measured the onset latency and peak width at half-maximum (PWHM); effects with onset latency >3.5 ms for spinal neurons or >5 ms for cortical neurons, and PWHM < 7 ms were identified as post-spike effects^1,8^.

The neurons that showed post-spike effects on at least one muscle were identified as PreM-INs or CM cells. If spinal neurons showed a large “motor unit” signature in the spike-triggered average of the unrectified EMG with only 50 spikes^11^, they were identified as putative motoneurons and excluded from the data set.

Mean percent increase of a post-spike effect was quantified by averaging the post-spike effect amplitude from the onset to offset, subtracting the baseline mean, and then dividing the result by the baseline mean and multiplying by 100. The muscle field was defined as a distribution of post-spike effects of a single PreM-IN or CM cell, and it was quantified as a vector of mean percent increase of post-spike effects.

### Temporal correlation between firing activity and target muscle activity

To quantify the temporal correlation between neuronal activity and target muscle activity, we calculated cross-correlations between the neuronal and muscle activities. First, the neuron instantaneous firing rate [*IFR*(*t*)] was calculated as the inverse of the interspike interval: *IFR*(*t*) = 1/(*t_i_*_+1_ − *t_i_*), for *t_i_* < *t* < *t_i_*_+1_, where *t_i_* is the time of the *i*th spike. The instantaneous firing rate was then low-pass filtered (second order, Butterworth, cutoff of 20 Hz in forward and backward directions) and down-sampled to 1000 Hz. Rectified EMGs were also low-pass filtered (second order, Butterworth, cutoff of 20 Hz in forward and backward directions) and down-sampled to 1000 Hz. Continuously recorded 90-s data points, which contained approximately 10 successive trials, were used to calculate the cross-correlation.

### Reconstruction of EMG signals from the post-spike effect of spinal interneurons

To estimate the contribution of single premotor neurons to the generation of muscle activity, we reconstructed the EMG signal by convoluting neural firing activity and a post-spike effect waveform. We used the spike-triggered average waveform, whose baseline trend was subtracted using the incremented-shifted averages method^6^, as a kernel. Because the waveform contained the EMG signal from 30 ms before to 50 ms after spike firing, we padded zeros for 20 ms before the waveform to centre the spike timing in the kernel. Next, we converted the spike timing signal to a continuous binary signal (0 or 1) at a 1-kHz sampling rate. Then, we convoluted the kernel with a spike count signal to reconstruct the EMG. Note that EMG signals were not normalized and were in voltage (µV) units to compare the original and reconstructed signals.

To evaluate the relative contribution of each premotor neuron, we averaged the original or reconstructed EMG signal, segmented from −1 to 3 s from grip onset, measured the area during the grip period (0 to 2 s from grip onset), and calculated the ratio of the area of the reconstructed EMG to that of the original EMG as an index of the relative contribution (% contribution).

### Extraction of muscle synergy

For the muscle synergy analysis, we selected 12 muscles which showed at least one post-spike effect by PreM-INs or CM cells (FDI, ADP, AbPB, AbDM, FDS, FDPr, FDPu, FCR, FCU, ED23, EDC, and ECU). Muscle synergies (synergy weights and synergy activity) were estimated from the EMGs using the non-negative matrix factorization (NMF)^12^. First, we removed electrical cross-talk between EMG signals by applying a blind-signal separation for the third-order differentiated EMG signals^10^. EMG signals were down-sampled to 1 kHz and differentiated three times. Then blind-signal separation was used with the preprocessed signals to remove the nonphysiological electrical cross-talk. After the separation, the signals were third-order integrated to restore the original waveforms.

The NMF was applied to the processed EMG signals. We used 8-min-long continuously recorded EMG signals. The separated EMGs were high-pass filtered (cutoff = 50 Hz), rectified, low-pass filtered (20 Hz), linear smoothed (100 time points), and down-sampled to 100 Hz. The amplitude of EMGs was normalized to set the mean amplitude to 1.

The NMF algorithm was initialized with random weight and activity matrices, which were drawn from a uniform distribution between 0 and 1. Values of these matrices were iteratively updated using the multiplicative rule until a change of EMG-reconstruction *R*^2^ < 0.01% in 20 consecutive iterations. Because the solutions for the synergies and their coefficients could fall into a local minimum, we repeated the synergy extraction 10 times from different initial values. We selected the synergies that showed the highest *R*^2^ for further analyses.

To identify the number of muscle synergies, we successively increased the number of synergies extracted from one to the number of muscles recorded (Extended Data Fig. 3). The *R*^2^ values were obtained with four-fold cross-validation by first extracting synergies from three of four 2-min-long data sets (the training sets) and then fitting the extracted synergies to the other unused quarter (the testing set). We averaged *R*^2^ values of the four-fold testing set and plotted them against the number of synergies extracted (Extended Data Fig. 3a–b). The number of synergies was defined as when *R*^2^ was closest to 0.9. The synergy weights and synergy activity were normalized to set the mean value of synergy activity to 1.

### Correlations of neural activity with muscle synergies and residuals

To compare the functional difference between PreM-INs and CM cells, we modelled hand muscle activity as a combination of lower and higher dimensional control, represented by muscle synergies and residuals, respectively. We calculated residuals by reconstructing the EMGs only with a linear combination of muscle synergies. As a definition, the muscle synergies account for approximately 90% of the variance of original EMGs. Then, we subtracted the reconstructed EMGs from the original EMGs to define residuals, which account for approximately 10% of the variance of the original EMGs.

The correlation of the temporal profiles of premotor neuronal activity with synergy activity or residuals was quantified with the response profile aligned to grip onset. Grip onset was defined as the time at which the rate of change of the total grip force exceeded 2 N/s. The response averages of instantaneous firing rate or synergy activity were compiled by aligning with the grip onset (from 1 s before to 3 s after grip onset) and averaging across trials. The correlation of the neural activity with a muscle synergy or a residual was quantified as *Pearson*’s correlation coefficient (*R*) between the two waveforms.

For each neuron–target muscle pair, we compared the neural correlation with the preferred muscle synergy and the residual of the target muscle. The preferred muscle synergy was defined with the similarity in the neuron’s muscle field and synergy weight, quantified as an absolute value of the cosine angle between the two vectors (muscle field and synergy weight) in muscle dimension (*n* = 12)^4^. This index ranged from 0 to 1, where 1 indicates perfect matching and 0 indicates no correlation. After we computed the similarity for all muscle synergies (three and four for monkeys E and S, respectively), we defined the muscle synergy with the highest similarity as a preferred synergy of the premotor neuron.

The preference index of each premotor neuron for the synergy or residual of the target muscle was calculated by subtracting the z-score correlation coefficient (atanh(*R*)) with the synergy from that of the residual (Fig. 4e). Therefore, a higher value indicates a higher preference for the residual, whereas a lower value indicates a higher preference for the synergy.

## Acknowledgements

We thank N. Takahashi, K. Takada, M. Togawa (National Institute for Physiological Sciences), and C. Sasaki (National Center of Neurology and Psychiatry) for technical assistance. We thank Lesley McCollum, PhD, from Edanz (https://jp.edanz.com/ac) for editing a draft of this manuscript. This work was supported by Grants-in-Aid (18020030, 18500315, 20020029, 23300143, 26120003, 26250013, 19H05724, 19H01092, 23H05488, K.S.; 06J02928, 21700437, 23700482, 19H03975, 19H05311, 21H00309, 22H04783, 24K02846 T.T.) from the Ministry of Education, Culture, Sports, Science and Technology of Japan (MEXT), the Japan Science and Technology Agency (JST) Precursory Research for Embryonic Science and Technology program (K.S.), JST FOREST Program (JPMJFR2045, T.T.), the JST Precursory Research for Embryonic Science and Technology program, and commissioned research (no. 22102) from the National Institute of Information and Communications Technology (NICT, K.S.), the Uehara Memorial Foundation, the Naito Foundation, the Takeda Science Foundation and the Brain Science Foundation (T.T.).

## Author Contributions

Conceptualization, T.T. and K.S.; Methodology, T.T., T.O., and K.S.; Investigation, T.T.; Writing – Original Draft, T.T. and K.S.; Writing – Review & Editing, T.T., T.O., and K.S.; Funding Acquisition, T.T. and K.S.

## Author information

Correspondence and requests for materials should be addressed to T.T. or K.S.

## Extended Data Figures

**Extended Data Figure 1.**
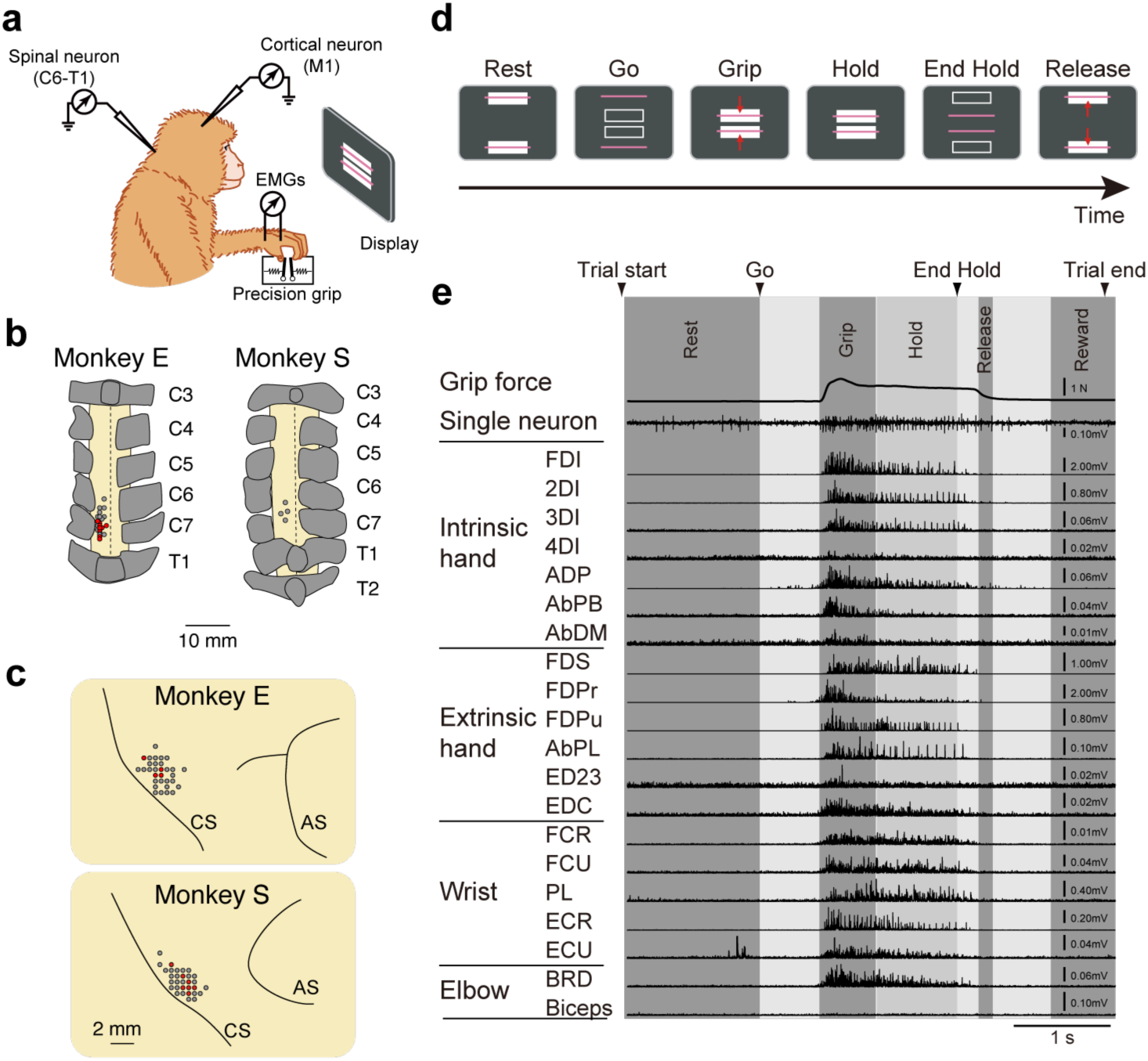
Experimental setups. **a**, Recording setup. We recorded single unit activity from spinal cervical cord or primary motor cortex and muscle activity of hand and arm muscles while the monkey was performing a precision grip task. **b,** Spinal recording sites in each monkey. Grey dots indicate the locations where we recorded the spinal neurons. Red dots indicate the location where we recorded the identified PreM-INs or CM cells. Letters on the right indicate the number of vertebrae. Dorsal views are shown. **c,** Cortical recording sites in each monkey. Grey and red dots in the same format as **b**. CS: central sulcus, AS: arcuate sulcus. **d**, Trial sequence. Monkeys were trained to grip, hold, and release the levers according to visual targets. **e**, Example of single-unit activity (PreM-IN), muscle activity (n = 20), and grip force.

**Extended Data Figure 2.**
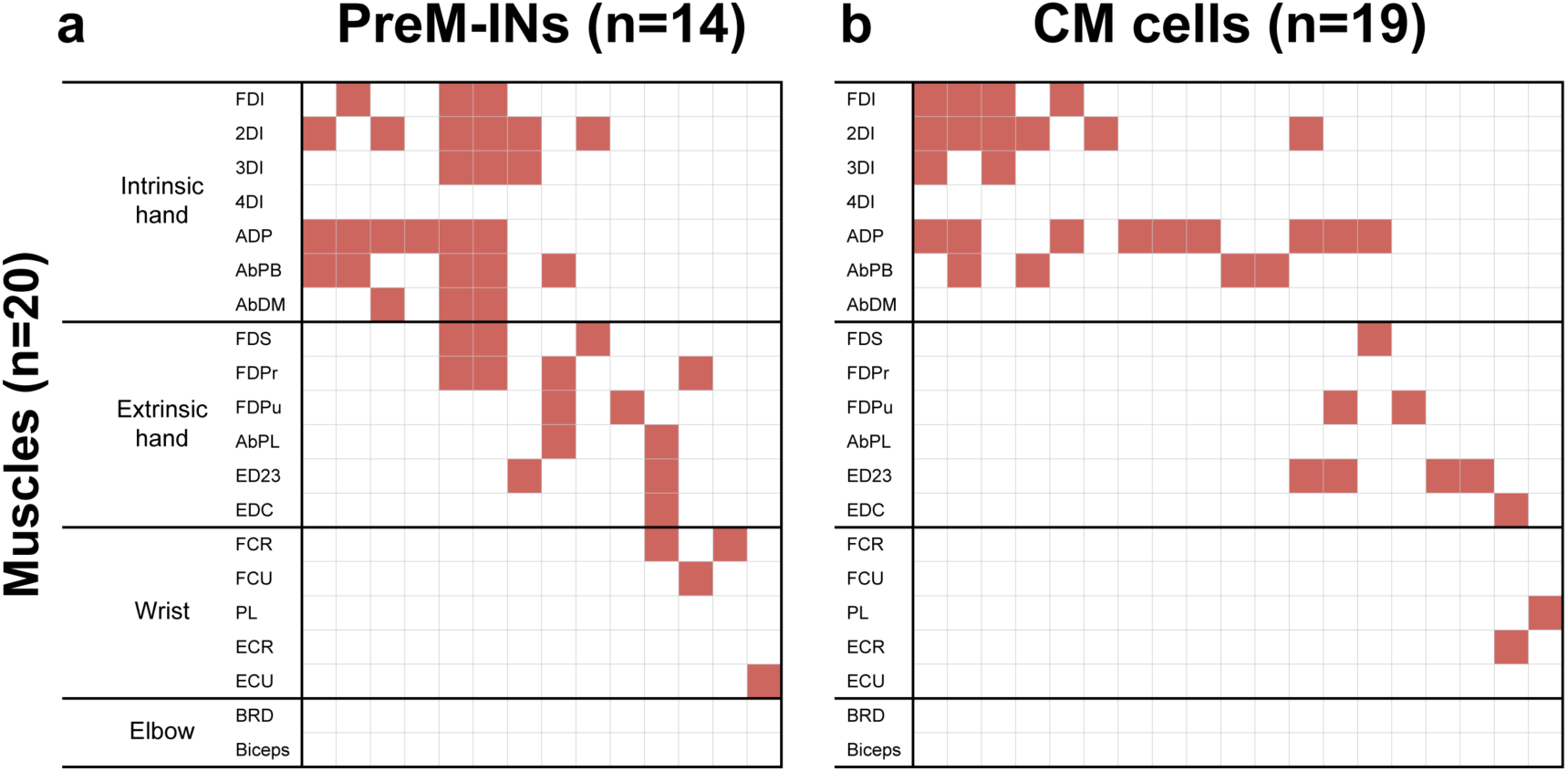
Distribution of post-spike facilitations of PreM-INs and CM cells. Post-spike facilitation produced by 14 PreM-INs (a) and 19 CM cells (b). Red squares indicate significant post-spike facilitations.

**Extended Data Figure 3.**
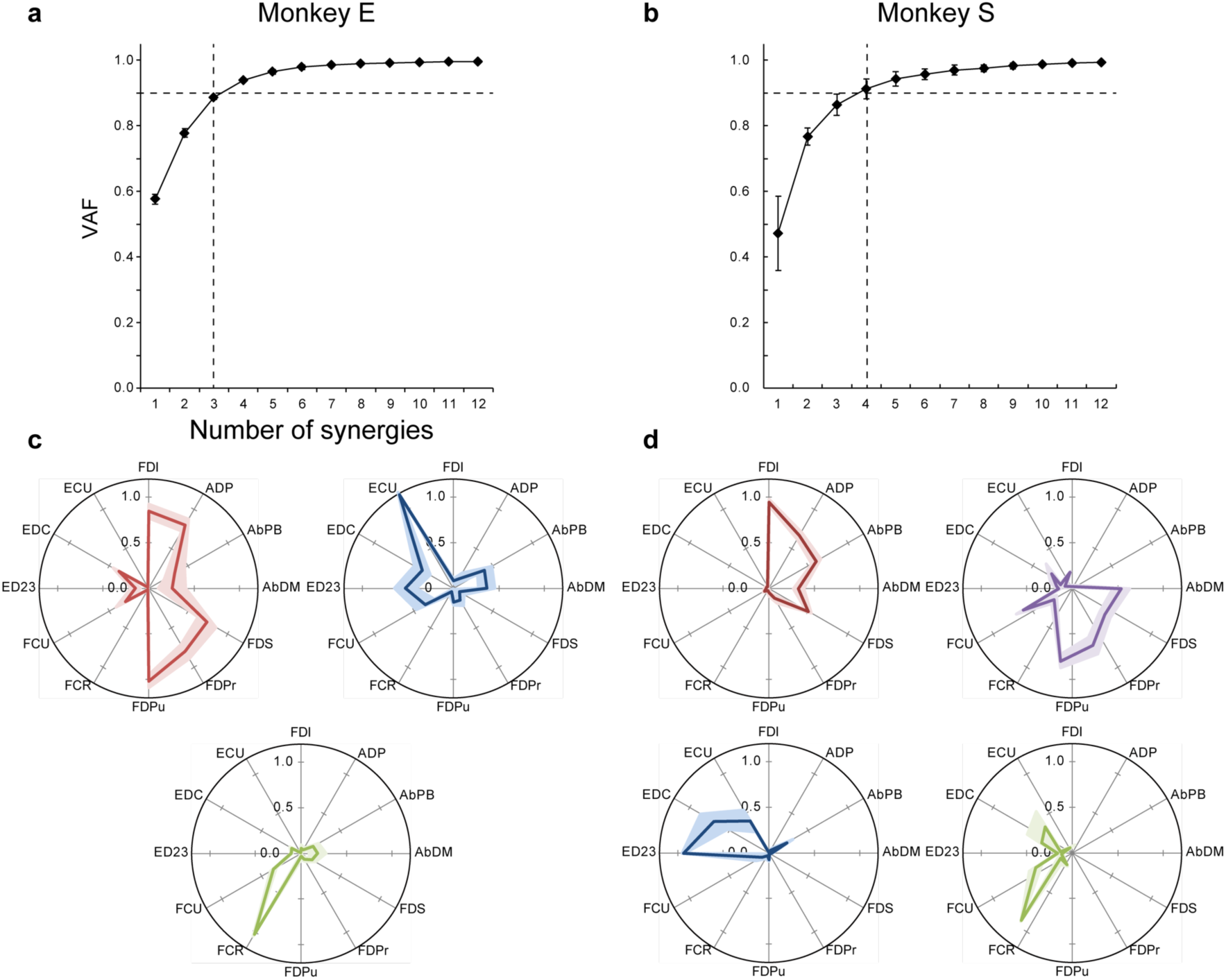
Extraction of muscle synergies. **a–b**, Selection of the number of muscle synergies. Variance accounted for (VAF) as a function of the number of extracted synergies for monkeys E (**a**) and S (**b**). The VAF curve was obtained by the fourfold cross-validation of the EMG data (filled circles). The number of muscle synergies (vertical dotted line) was selected where the VAF was closest to 0.9 (horizontal dotted line). Error bars indicate SEM across sessions. **c–d**, Muscle synergies extracted from monkeys E (**c**) and S (**d**).

**Extended Data Figure 4.**
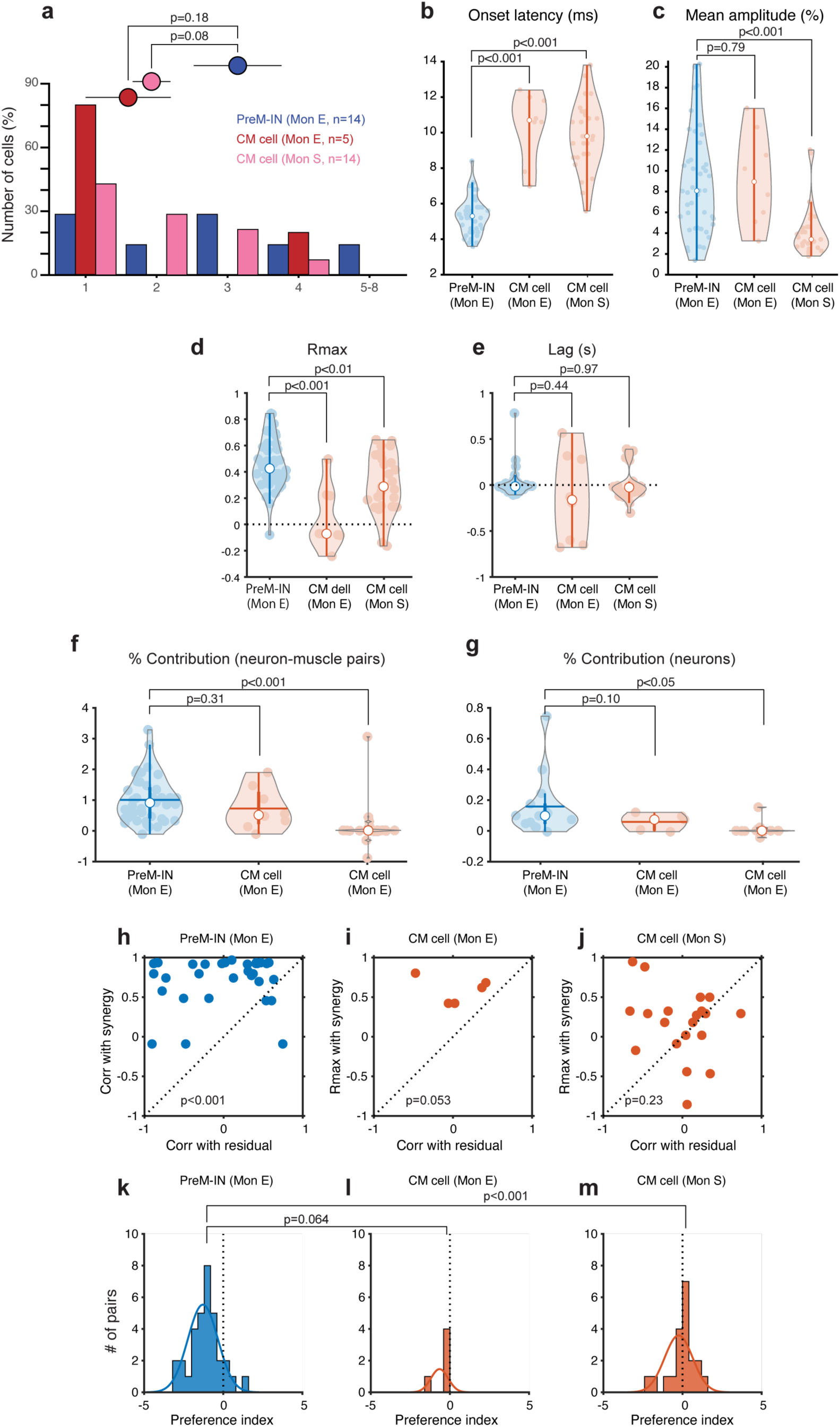
Results separated for each animal. Data were compared between PreM-INs and CM cells of monkey E or that of monkey S. **a,** Histogram of muscle field size. Same format as Fig. 1b. **b–c,** Onset latency and mean amplitude of post-spike facilitations. Same format as Fig. 1c–d. **d–e,** *Rmax* and lag of temporal correlation. Same format as Fig. 2g–h. **f–g,** Percent contribution of neuron– muscle pairs and neurons. Same format as Fig. 3d–e. **h–j,** Correlation with the preferred synergy and residual of the target muscle. Same format as Fig. 4d. **k–m,** Preference index. Same format as Fig. 4d.

## References

1. Porter, R. & Lemon, R. Corticospinal Function and Voluntary Movement. (Oxford University Press, 1993).

2. Lemon, R. N. Descending pathways in motor control. Annual review of neuroscience 31, 195–218 (2008).

3. Rathelot, J.-A. & Strick, P. L. Subdivisions of primary motor cortex based on cortico-motoneuronal cells. Proceedings of the National Academy of Sciences of the United States of America 106, 918–923 (2009).

4. Strick, P. L., Dum, R. P. & Rathelot, J.-A. The Cortical Motor Areas and the Emergence of Motor Skills: A Neuroanatomical Perspective. Annu Rev Neurosci 44, 1– 23 (2021).

5. Takei, T. & Seki, K. Spinal Interneurons Facilitate Coactivation of Hand Muscles during a Precision Grip Task in Monkeys. J. Neurosci. 30, 17041–17050 (2010).

6. Takei, T. & Seki, K. Spinal Premotor Interneurons Mediate Dynamic and Static Motor Commands for Precision Grip in Monkeys. J. Neurosci. 33, 8850–8860 (2013).

7. Takei, T. & Seki, K. Synaptic and functional linkages between spinal premotor interneurons and hand-muscle activity during precision grip. Front. Comput. Neurosci. 7, 40 (2013).

8. Fetz, E. E. & Cheney, P. D. Postspike facilitation of forelimb muscle activity by primate corticomotoneuronal cells. Journal of neurophysiology 44, 751–772 (1980).

9. Buys, E. J., Lemon, R. N., Mantel, G. W. & Muir, R. B. Selective facilitation of different hand muscles by single corticospinal neurones in the conscious monkey. The Journal of physiology 381, 529–549 (1986).

10. Takei, T., Confais, J., Tomatsu, S., Oya, T. & Seki, K. Neural basis for hand muscle synergies in the primate spinal cord. Proc. Natl. Acad. Sci. 114, 8643–8648 (2017).

11. Yan, Y., Goodman, J. M., Moore, D. D., Solla, S. A. & Bensmaia, S. J. Unexpected complexity of everyday manual behaviors. Nat Commun 11, 3564 (2020).

12. Santello, M., Flanders, M. & Soechting, J. F. Postural hand synergies for tool use. The Journal of neuroscience: the official journal of the Society for Neuroscience 18, 10105–10115 (1998).

13. Marshall, N. J. et al. Flexible neural control of motor units. Nat Neurosci 1–13 (2022) doi:10.1038/s41593-022-01165-8.

14. Bizzi, E., Cheung, V. C. K., d’Avella, A., Saltiel, P. & Tresch, M. Combining modules for movement. Brain research reviews 57, 125–133 (2008).

15. Giszter, S. F. Motor primitives-new data and future questions. Current opinion in neurobiology 33, 156–165 (2015).

16. Berniker, M., Jarc, A., Bizzi, E. & Tresch, M. C. Simplified and effective motor control based on muscle synergies to exploit musculoskeletal dynamics. Proceedings of the National Academy of Sciences of the United States of America 106, 7601–7606 (2009).

17. Cheung, V. C. K. et al. Stability of muscle synergies for voluntary actions after cortical stroke in humans. Proceedings of the National Academy of Sciences of the United States of America 106, 19563–19568 (2009).

18. Song, Y., Hirashima, M. & Takei, T. Neural Network Models for Spinal Implementation of Muscle Synergies. Front. Syst. Neurosci. 16, 800628 (2022).

19. Ting, L. H. & McKay, J. L. Neuromechanics of muscle synergies for posture and movement. Current opinion in neurobiology 17, 622–628 (2007).

20. Bennett, K. M. & Lemon, R. N. Corticomotoneuronal contribution to the fractionation of muscle activity during precision grip in the monkey. J Neurophysiol 75, 1826–1842 (1996).

21. Schieber, M. H. Individuated finger movements of rhesus monkeys: a means of quantifying the independence of the digits. J Neurophysiol 65, 1381–1391 (1991).

22. Schieber, M. H. Muscular production of individuated finger movements: the roles of extrinsic finger muscles. The Journal of neuroscience: the official journal of the Society for Neuroscience 15, 284–297 (1995).

23. Muir, R. B. & Lemon, R. N. Corticospinal neurons with a special role in precision grip. Brain Res 261, 312–316 (1983).

24. Lemon, R. N., Mantel, G. W. & Muir, R. B. Corticospinal facilitation of hand muscles during voluntary movement in the conscious monkey. The Journal of physiology 381, 497–527 (1986).

25. Maier, M. A., Bennett, K. M., Hepp-Reymond, M. C. & Lemon, R. N. Contribution of the monkey corticomotoneuronal system to the control of force in precision grip. Journal of neurophysiology 69, 772–785 (1993).

26. Schieber, M. H. & Rivlis, G. Partial reconstruction of muscle activity from a pruned network of diverse motor cortex neurons. Journal of neurophysiology 97, 70–82 (2007).

27. Schieber, M. H. & Rivlis, G. A spectrum from pure post-spike effects to synchrony effects in spike-triggered averages of electromyographic activity during skilled finger movements. Journal of neurophysiology 94, 3325–3341 (2005).

28. Poliakov, A. V. & Schieber, M. H. Multiple fragment statistical analysis of post-spike effects in spike-triggered averages of rectified EMG. Journal of neuroscience methods 79, 143–150 (1998).

29. Griffin, D. M., Hoffman, D. S. & Strick, P. L. Corticomotoneuronal cells are “functionally tuned.” Science (New York, NY) 350, 667–670 (2015).

30. Cheney, P. D. & Fetz, E. E. Functional classes of primate corticomotoneuronal cells and their relation to active force. Journal of neurophysiology 44, 773–791 (1980).

31. Armand, J., Olivier, E., Edgley, S. A. & Lemon, R. N. Postnatal development of corticospinal projections from motor cortex to the cervical enlargement in the macaque monkey. The Journal of neuroscience: the official journal of the Society for Neuroscience 17, 251–266 (1997).

32. Perlmutter, S. I., Maier, M. A. & Fetz, E. E. Activity of spinal interneurons and their effects on forearm muscles during voluntary wrist movements in the monkey. Journal of neurophysiology 80, 2475–2494 (1998).

33. Long, C., Conrad, P. W., Hall, E. A. & Furler, S. L. Intrinsic-extrinsic muscle control of the hand in power grip and precision handling. An electromyographic study. The Journal of bone and joint surgery American volume 52, 853–867 (1970).

34. Smith, A. M. The coactivation of antagonist muscles. Canadian journal of physiology and pharmacology 59, 733–747 (1981).

35. Maier, M. A. & Hepp-Reymond, M. C. EMG activation patterns during force production in precision grip. I. Contribution of 15 finger muscles to isometric force. Experimental brain research 103, 108–122 (1995).

36. Kasser, R. J. & Cheney, P. D. Characteristics of corticomotoneuronal postspike facilitation and reciprocal suppression of EMG activity in the monkey. J Neurophysiol 53, 959–978 (1985).

37. Fetz, E. E., Perlmutter, S. I., Prut, Y., Seki, K. & Votaw, S. Roles of primate spinal interneurons in preparation and execution of voluntary hand movement. Brain Res. Rev. 40, 53–65 (2002).

38. Baldissera, F., Hultborn, H. & Illert, M. Integration in spinal neuronal systems. in 509–595 (1981). doi:10.1002/cphy.cp010212.

## Additional references

1. Takei, T. & Seki, K. Spinal Interneurons Facilitate Coactivation of Hand Muscles during a Precision Grip Task in Monkeys. J. Neurosci. 30, 17041–17050 (2010).

2. Takei, T. & Seki, K. Synaptic and functional linkages between spinal premotor interneurons and hand-muscle activity during precision grip. Front. Comput. Neurosci. 7, 40 (2013).

3. Takei, T. & Seki, K. Spinal Premotor Interneurons Mediate Dynamic and Static Motor Commands for Precision Grip in Monkeys. J. Neurosci. 33, 8850–8860 (2013).

4. Takei, T., Confais, J., Tomatsu, S., Oya, T. & Seki, K. Neural basis for hand muscle synergies in the primate spinal cord. Proc. Natl. Acad. Sci. 114, 8643–8648 (2017).

5. Takei, T. & Seki, K. Spinomuscular Coherence in Monkeys Performing a Precision Grip Task. J. Neurophysiol. 99, 2012–2020 (2008).

6. Davidson, A. G., O’Dell, R., Chan, V. & Schieber, M. H. Comparing effects in spike-triggered averages of rectified EMG across different behaviors. Journal of neuroscience methods 163, 283–294 (2007).

7. Poliakov, A. V. & Schieber, M. H. Multiple fragment statistical analysis of post-spike effects in spike-triggered averages of rectified EMG. Journal of neuroscience methods 79, 143–150 (1998).

8. Schieber, M. H. & Rivlis, G. A spectrum from pure post-spike effects to synchrony effects in spike-triggered averages of electromyographic activity during skilled finger movements. Journal of neurophysiology 94, 3325–3341 (2005).

10. Kilner, J. M., Baker, S. N. & Lemon, R. N. A novel algorithm to remove electrical cross-talk between surface EMG recordings and its application to the measurement of short-term synchronisation in humans. J Physiology 538, 919–930 (2002).

11. Maier, M. A., Perlmutter, S. I. & Fetz, E. E. Response patterns and force relations of monkey spinal interneurons during active wrist movement. Journal of neurophysiology 80, 2495–2513 (1998).

12. Lee, D. D. & Seung, H. S. Learning the parts of objects by non-negative matrix factorization. Nature 401, 788–791 (1999).

